# Quantification of fast molecular adhesion by fluorescence footprinting

**DOI:** 10.1101/2021.01.05.425475

**Authors:** Adam B. Yasunaga, Isaac T.S. Li

## Abstract

Rolling adhesion is a unique process in which the adhesion events are short-lived and operate under highly non-equilibrium conditions. These characteristics pose a challenge in molecular force quantification, where *in situ* measurement of such forces cannot be achieved with most molecular force sensors that probe near equilibrium. In this report, we demonstrated a quantitative adhesion footprint assay combining DNA-based non-equilibrium force probes and modelling to measure the molecular force involved in fast rolling adhesion. We were able to directly profile the ensemble molecular force distribution during rolling adhesion with a dynamic range between 0 – 18 pN. Our results showed that the shear stress driving bead rolling motility directly controls the molecular tension on the probe-conjugated adhesion complex. Furthermore, the shear stress can steer the dissociation bias of components within the molecular force probe complex, favouring either DNA probe dissociation or receptor-ligand dissociation.

## Introduction

The rolling adhesion cascade is a crucial phenomenon in the immune system where leukocytes rapidly roll on the blood vessel walls to reach the site of infection (*1, 2*). Rolling adhesion is mediated by the interaction between P-selectin on the surface of endothelial cells, and P-selectin glycoprotein ligand-1 (PSGL1) on the surface of leukocytes (*2, 3*). Selectins exhibit both a high on and off rate with high affinity allowing for rapid formation and dissociation of P-selectin/PSGL1 interactions during rolling (*4*–*6*). This interaction has also been shown to display a catch-slip bond response to tensile force, where bond lifetime initially increases (catch bond) with force followed by a decrease in lifetime (slip bond) after a threshold force has been crossed (*2, 7, 8*). In addition to rolling, PSGL1 acts as a mechanosensitive receptor that, upon engagement with P-selectin, triggers the subsequent firm adhesion stage of the rolling adhesion cascade (*1, 3, 9*). With force playing a role in both adhesion and signaling, it is imperative to understand the dynamics of tensile force on the P-selectin/PSGL1 interaction under physiological rolling conditions. However, no direct experimental measurement of these forces is available due to challenges in quantifying forces in such a dynamic system, where individual adhesion bonds are only briefly (millisecond time scale) subjected to tension once.

Although the forces on the P-selectin/PSGL1 interaction have not been directly investigated during cell rolling, the force response of the interaction has been studied *in vitro* at the single-molecule level (*10*). This has been accomplished using atomic force microscopy (AFM) (*4, 7, 11, 12*), biomembrane force probe (BFP) (*13*) and optical tweezers (OT) (*10, 14*). In each of these approaches, the force response of the interaction is studied by bringing a receptor-coated surface and a ligand-coated surface together and then retracting one surface to apply a tensile force on the existing interaction(s). Depending on the technique used to probe the P-selectin/PSGL1 interaction, the rupture forces have been reported to be between 0.08 and 250pN (*10*). However, without direct measurement in a real cell rolling system, it is unknown what force range reported by in vitro measurements are physiologically relevant. In addition, the complex adhesion tether geometry and cell rolling dynamics makes it difficult to accurately replicate the force loading history by instrument-based force-spectroscopy techniques. Therefore, an experimental approach to directly measure molecular adhesion forces in a rolling adhesion system is needed. Although currently existing molecular force sensors have been deployed successfully in measuring tension in cell adhesion involving integrins (*15, 16*), T-cell receptors (*17, 18*), cadherins (*19*–*21*), and cytoskeletal proteins (*22*), these adhesion events occur on a time-scale (minutes) (*23*) much slower comparing to those involving selectin-family proteins in cell rolling (millisecond) (*6, 23, 24*). Because of the fast dynamics of these adhesion interactions, there is no currently available molecular force quantification technique compatible with rolling adhesion. This is a fundamental challenge in molecular and cellular force quantification, as selectins are one of the major classes of cell adhesion molecules (CAMs), and accurate force measurement and control have profound implication in understanding their coupled signalling pathways in disease pathophysiology (*1, 9, 25*).

Current molecular force sensors (MFS) fall into three general categories based on their force sensing mechanisms, as reviewed previously (*26*). These categories are defined as reversible analog, reversible digital, and irreversible sensors. Reversible MFSs are not suitable for studying rolling adhesion because the signal-to-noise ratio is insufficient to measure rapid, non-equilibrium, force changes during the short selectin-mediated bond formation and dissociation at the surface density required to sustain rolling adhesion. Irreversible MFSs, on the other hand, are well-suited for non-equilibrium processes as they can produce a permanent record of a single adhesion event with significantly higher signal-to-noise ratio. These desirable properties led to the development of the adhesion footprint assay (*27*) based on tension gauge tether (TGT) (*28*), a DNA-based non-equilibrium force sensor, where the spatial distribution of rolling adhesion was recorded and mapped. However, quantification of adhesion force was limited, as the fluorescence intensity in an “adhesion footprint” only represents the number of adhesion events rupturing the TGT, and does not quantify the molecular forces on the P-selectin/PSGL1 interactions (*26*). In this article, we developed a framework to quantify the adhesion forces using adhesion footprint assay. Using a bead rolling adhesion model system, we obtained the distribution of instantaneous molecular adhesion force and rupture force of the P-selectin/PSGL1 interaction during rolling adhesion.

## Results

### Bead rolling adhesion as a model system

To investigate the forces involved in rolling adhesion, we used bead rolling as our model system. This was accomplished by rolling PSGL1 coated polystyrene beads on a surface functionalized with P-selectin conjugated TGTs (Fig. 1A). By using beads in place of live cells, we had a consistent and well-defined system allowing us to focus solely on the molecular forces on the P-selectin/PSGL1 interactions. Unlike with cells, beads allowed us to make the assumptions that the shape of the beads remains the same regardless of the applied force, and there is a uniform distribution of PSGL1 receptors on the bead surface. With these assumptions, we were able to develop a model of bead rolling adhesion that is critical to the association of the “adhesion footprint” fluorescence intensity to molecular force on the P-selectin/PSGL1 interactions.

**Fig. 1.**
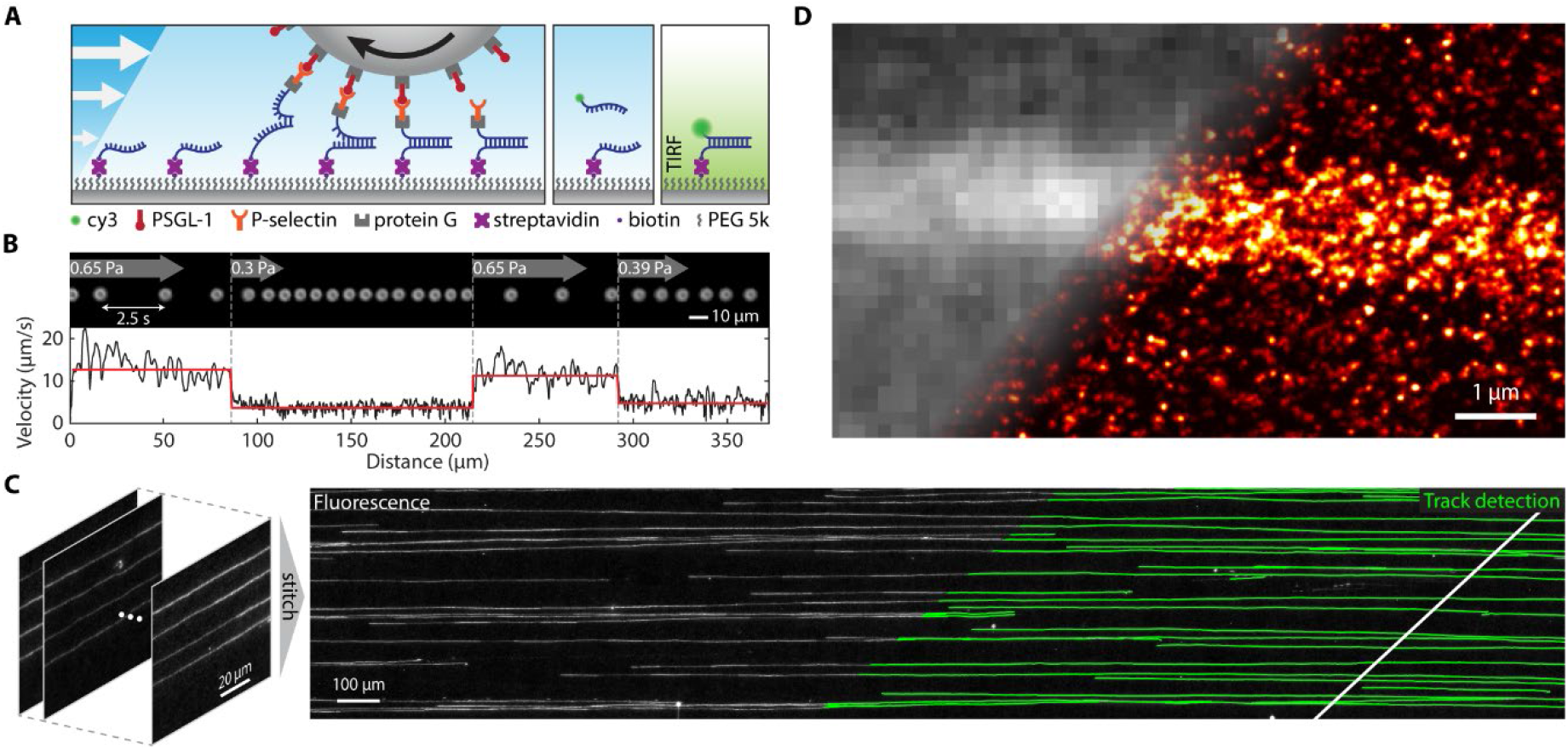
Bead rolling adhesion with adhesion footprint assay. **(A)** Schematic of the rolling adhesion footprint assay. A PSGL1 coated bead rolls on a surface by binding to P-selectin and eventually rupturing the conjugated DNA duplex, leaving a single stranded DNA (ssDNA) on the surface corresponding to the precise locations of each adhesion event. This adhesion footprint assay leaves a trail of ssDNA on the surface that can be fluorescently labeled with a complementary strand and further imaged with TIRF microscopy to observe the tracks left by the rolling beads. **(B)** Darkfield imaging of bead rolling under altering steps of shear stress. The top image imposes snapshots of a bead every 2.5 seconds in response to the different shear stresses. The bottom panel shows the corresponding instantaneous velocity (black) and mean velocity (red). **(C)**. A stitched image showing the fluorescence adhesion footprint tracks (left) that are observed after PSGL1 beads roll on a P-selectin/TGT surface. Individual tracks (green lines) are detected (right), enabling isolation and analysis of the fluorescence intensity trajectory of individual beads. **(D)**. The diffraction limited and DNA PAINT super-resolution image of a single fluorescence track.

The bead rolling assay was performed by flowing PSGL1 coated beads through a parallel plate flow chamber functionalized by TGTs conjugated to P-selectin. The shear stress in the chamber was controlled by a syringe pump, allowing for a user-defined sequence of shear stresses to be applied. A darkfield microscope was used to record and track the rolling beads (Fig. 1B) and an inverted total internal reflection fluorescence (TIRF) microscope was used to image the fluorescence tracks left on the surface by the beads (Fig. 1A, C). We hypothesized that increasing shear stress on a rolling bead leads to an increase in molecular adhesion forces, shorter adhesion bond lifetime, and faster rolling velocity. Therefore, in this study, beads were subjected to various physiological shear stresses to probe the physiological P-selectin/PSGL1 adhesion force during rolling adhesion. At each shear stress, both velocity and fluorescence intensity data were analyzed and used in conjunction with a model of bead rolling adhesion to quantify the molecular forces on the P-selectin/PSGL1 interaction.

By doing single particle tracking analysis of darkfield bead rolling movies, we showed that not only does the bead roll on the P-selectin/TGT surface, the rolling adhesion is stable, as seen from the uniform rolling velocity of a single bead (Fig. 1A). While it is possible to track a single bead over multiple steps of shear stresses under 10x magnification, doing so at 100x magnification under TIRF imaging condition is challenging. Hence, instead of live tracking, we scanned the surface and stitched 2000-3000 individual TIRF images to form a large image composed of many tracks, each over multiple cycles of shear stress steps (Fig. 1C). Individual tracks were detected by custom-written code, and isolated for subsequent analysis.

The adhesion footprint assay is intrinsically compatible with super-resolution DNA PAINT, as the DNA unzipping leaves a single stranded DNA on the surface that can be imaged directly by DNA PAINT imaging strands. Doing so allowed us to directly visualize individual ruptured TGT on the surface (Fig. 1D). The density of ruptured TGT in a typical track in our experiment ranges between ∼10 to 40 per µm^2^ (excluding background) while the total surface density of TGT is estimated to be ∼ 2000 per µm^2^. This is a clear demonstration of the high signal-to-noise ratio of TGT-based adhesion footprint assay and the necessity of this approach for fast cell adhesion studies. Given the rapid motility rolling bead/cell, the receptors have little time to form tether with ligands. Hence, even though both surfaces have saturating density of receptors and ligands, only 0.5% to 2.0% of all TGTs form tethers with the bead to get ruptured. To distinguish the signal and quantify forces on such a small proportion of tethered force sensors among a large background of non-tethered force sensors would pose a significant detection challenge for force sensors measuring equilibrium forces using FRET or fluorescence quenching.

### Shear stress controls bead rolling velocity and track fluorescence intensity

Shear flow enables the rolling adhesion by applying an overall force to the bead, which redistributes among adhesion tethers. It is the stochastic breaking of these tethers that enables a bead to roll in the direction of the flow. The observed rolling velocity is directly related to the force on the adhesive interactions during rolling adhesion; and by changing the shear stress on the beads, we can effectively change the force on the P-selectin/PSGL1 interactions. Greater shear stress translates directly to greater tension among the adhesion tethers. In an effort to understand the molecular force on catch-bonds in a real rolling system under physiological condition, we changed shear stress to put the system in force regimes where either the catch-bond or the slip-bond behaviours dominated.

First, we track the rolling velocity of individual beads to understand its response to shear stress. A syringe pump was programmed to apply a custom shear flow profile (Fig. 2A). Instantaneous rolling velocity (Fig. 2B) of an individual bead was determined by tracking its centroid using a custom-written MATLAB code. Our analysis showed that bead rolling velocity scales exponentially to shear stress at both the single bead and population levels (Fig. S1). This relationship is primarily dictated by the force-dependent dissociation of either TGT or P-selectin/PSGL1 within the complex and exhibits an apparent slip-bond behaviour. The mean velocity (*v*) of a steadily rolling bead is inversely proportional to the mean life-time of the adhesion complex due to simple geometry, while the force-dependent life-time (*τ*_*mol*_) of a slip bond is characterized by the following exponential function, where τ^0^_mol_ is the zero-force lifetime, F is the molecular tension, *x*^*‡*^ is the distance to dissociation barrier:

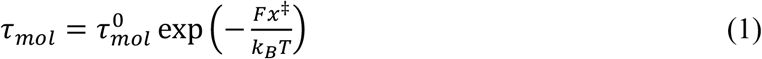

Given that the shear stress (*τ*_*shear*_) is directly proportional to the molecular forces on individual adhesion bonds under steady-state rolling, it follows that the mean rolling velocity is an exponential function of shear stress, where c_1_ and c_2_ are constants:

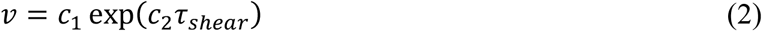

The agreement between this simple explanation and the experimental result on the exponential nature of *v* vs. *τ*_*shear*_ seems suggest that the rolling adhesion system here operates primarily in a regime where the force-dependent life-time of slip-bonds dominates. However, no direct relation between the behaviour and the molecular forces in these systems can be established, even though whole cell/bead tracking has been traditionally used to infer what is happening at the molecular scale. This calls for a direct measurement of the molecular forces involved in rolling adhesion.

**Fig. 2.**
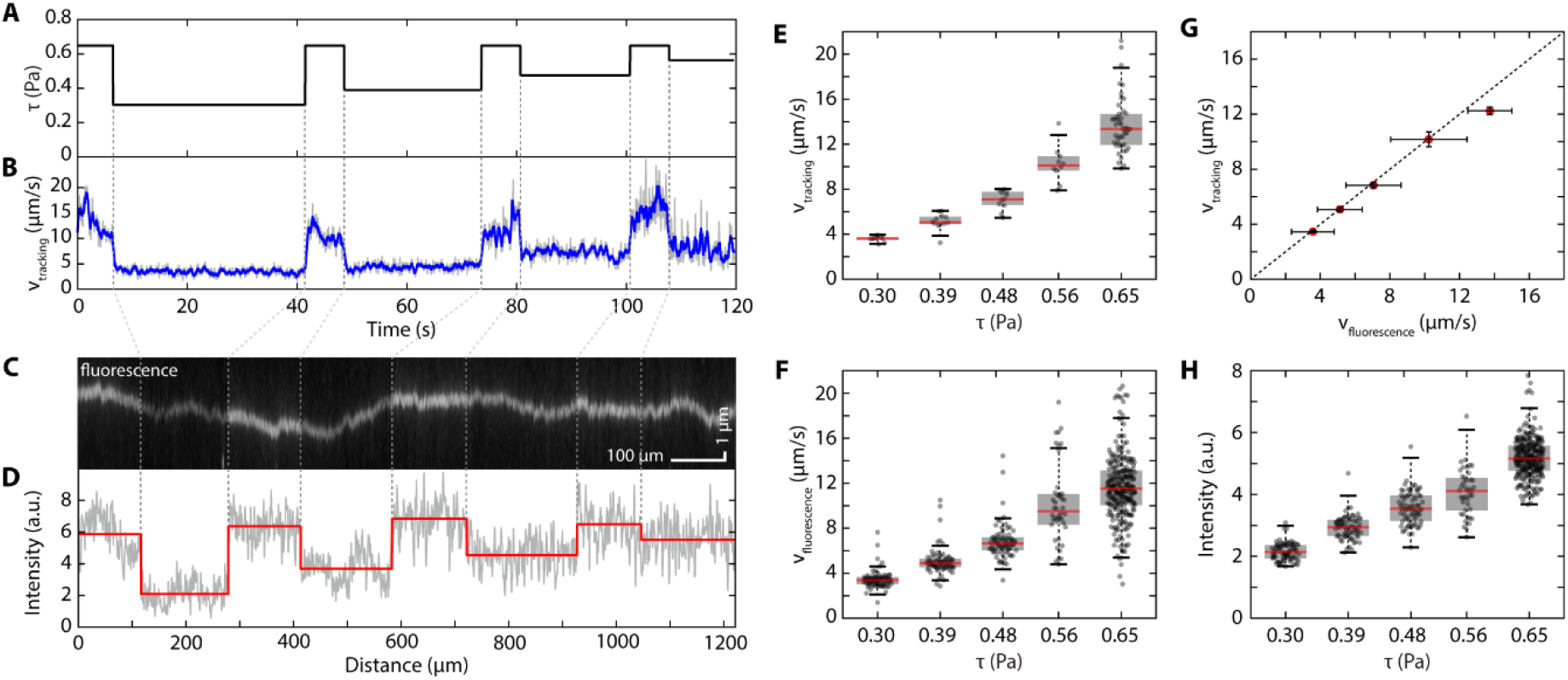
Shear stress dictates bead rolling velocity and fluorescence track intensity. **(A)** Syringe pump-controlled flow series applied to the rolling beads. **(B)** Population mean rolling velocity of PSGL1 coated beads over time resulting from the applied flow series shown in (A). **(C)** Representative fluorescence track imaged with TIRF after rolling beads on the TGT/P-selectin surface with the flow series show in (A). **(D)** Intensity trace along the length of the track showing raw data in gray and the mean intensity at each shear stress from (A) in red. **(E)** Mean tracking bead velocity as a function of shear stress. Tracking bead velocity was determined through single particle tracking of rolling beads. **(F)** Mean fluorescence bead velocity as a function of shear stress. Fluorescence bead velocity was determined by the length of each segment seen in a fluorescence track and the duration corresponding the segments. **(G)** Scatter plot of fluorescence velocity vs tracking bead velocity showing a positive correlation (r = 0.99, p < 0.001) with error bars representing standard error of the mean. **(H)** Fluorescence track intensity as a function of shear stress.

Unlike single molecule studies where force-extension on a single adhesion bond can be directly measured, at any given time, many adhesion bonds stretched to different tension are simultaneously involved in rolling adhesion. This makes it challenging to directly measure the magnitude of forces on individual molecules. Instead, our method focuses on measuring the distribution of molecular forces, which provides meaningful insight on the force evolution history of individual bonds during rolling. To investigate this distribution, we applied the adhesion footprint assay (*27*) (Fig. 1A) where P-selectin functionalized TGTs were used to fluorescently report and quantify molecular adhesion forces. The differential force-dependent dissociation kinetics of P-selectin/PSGL1 interaction vs. dsDNA unzipping leads to dissociation probabilities biased towards either P-selectin/PSGL1 (under low force), or dsDNA unzipping (under high force). Therefore, a single bead rolling on P-selectin functionalized TGTs at different shear stress will produce a trail of ruptured TGTs behind it (Fig. 1A), with densities dependent on the shear stress. Following each bead rolling experiment, the ruptured TGTs were fluorescently labeled (Fig. 1A) to reveal the tracks left on the surface by the rolling beads (Fig. 1C).

The intensity of these tracks is directly proportional to the number of ruptured TGTs and hence will lead us to dissecting the molecular force distribution. The fluorescence tracks were imaged with TIRF microscopy and each track was isolated and analyzed with custom-written MATLAB code. Upon observing the fluorescence tracks, it was seen that each track had a pattern of bright and dim segments along the length of the tracks (Fig. 2C). Similar to rolling velocity, the fluorescence intensity along the length of each track (Fig. 2D) also mimicked the pattern of the flow series used to roll the beads. Because bead rolling movies are collected at a different magnification on a different microscope, they cannot be directly correlated to the fluorescence tracks. In order to investigate the relationships between fluorescence intensity, bead velocity, and shear stress of a single bead, this information needs to be derived directly from individual fluorescence tracks. To do so, each track was split into segments, each corresponding to a single shear stress (Fig. 2A) where the mean fluorescence intensity (Fig. 2F) was calculated. In addition, the mean rolling velocity was calculated for each segment based on segment length and the duration for which the corresponding shear stress was applied. In doing this, a second set of velocity data was generated strictly from the fluorescence track data. The single-bead velocity distribution from fluorescent track analysis agrees with the single-bead velocity distribution from tracking the bead movie (Fig. 2G) with a strong positive correlation (R = 0.99, p < 0.001). Therefore, the velocity data extracted from the fluorescence tracks was accurate and was used for all subsequent analysis.

Analysis of the fluorescence tracks showed that track intensity increases monotonically to shear stress (Fig. 2H). As the fluorescence intensity is dependent on the surface density of ruptured TGTs, contrast in the fluorescence intensity indicates that our method is sensitive enough to molecular force changes within range of our shear stresses. Given our previous work, higher molecular force corresponds to greater amount of ruptured TGT, leading to greater fluorescence intensity. Hence, the result is consistent with our expectation. However, fluorescence intensity only shows the relative overall change in molecular force. It does not provide a direct quantification of the distribution of molecular forces. Therefore, a bead rolling adhesion model taking into account the force-dependent dissociation behaviour of serially connected TGT and P-selectin/PSGL1 is needed to quantify the molecular forces involved in rolling adhesion.

### Modelling bead rolling in the adhesion footprint assay

In order to determine the magnitude of the molecular force from the experimentally observed fluorescence intensity, we developed a robust model to describe the molecular force distribution during bead rolling adhesion. The model assumes that a hard-sphere rolls at a constant velocity (Fig. 3A), which is consistent with our experimental observation (Fig. 2B). Without solving Newtonian mechanics, this assumption leads to a steady-state condition that allows us to numerically solve the evolution of molecular force across individual adhesion tethers. Under this steady-state condition, every molecular tether follows the same end-to-end extension profile over time, *d(t)* as defined entirely by geometry (Fig. 3A):

**Fig. 3.**
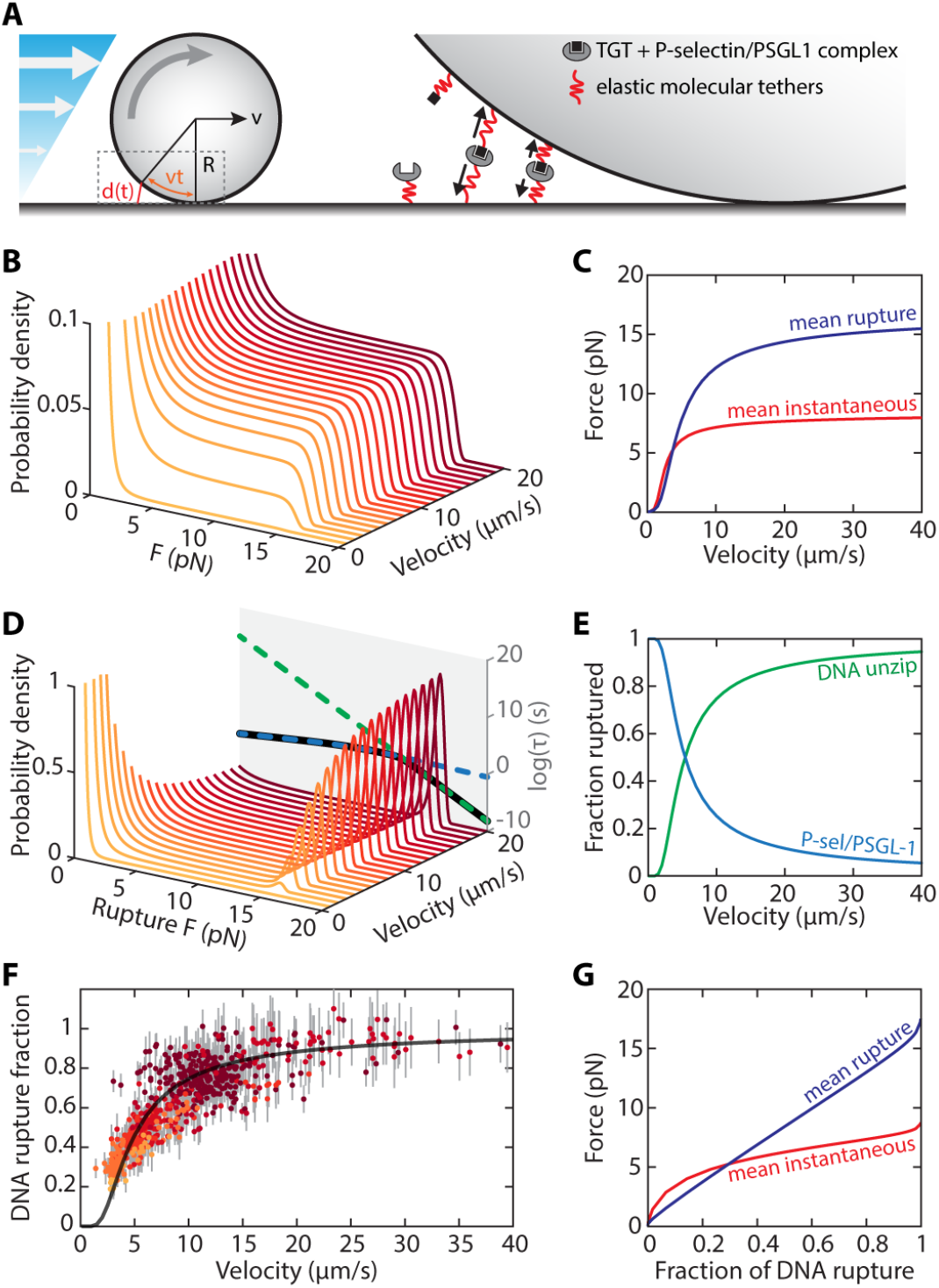
Modelling force distribution in the adhesion footprint assay. **(A)** The steady-state bead rolling model with parameters indicated in the schematic. The zoomed-in picture shows the molecular tethers are being stretched and dissociate as the bead roll forward, mimicking a force-extension experiment that acts on many tethers. **(B)** Instantaneous force distribution at different bead rolling velocity. **(C)** The expected instantaneous (red) and rupture (blue) force as a function of bead velocity. **(D)** Rupture force distribution at different bead rolling velocity. The force-dependent life-time of P-selectin/PSGL1 (blue dash), DNA reporter unzipping (green dash), and the overall bond rupture profile (black). **(E)** The fraction of bond rupture due to DNA unzipping and P-selectin/PSGL1 dissociation at different bead velocities. **(F)** Normalized, track fluorescence intensity as a function of velocity of individual beads, each data point is coloured by the shear stress it experiences under. Error bars indicate standard deviation of the fluorescence intensity along each track. Solid black line is the calculated DNA rupture fraction as a function of bead velocity from the model. **(G)** The expected rupture (blue) and instantaneous (red) force as a function of the DNA rupture fraction, predicted by the model. This is essentially a force-fluorescence calibration curve.

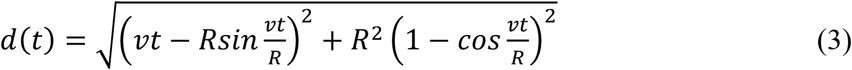

where *R* is the radius of the bead and *v* is the rolling velocity. Hence, the force loading history *f(t)* of each adhesion bond can be determined by the numerically evaluating the force-extension profile *f(d)* of the molecular tether. Individual components of the molecular tether used in our experiment was modelled using worm-like-chain parameters from the literature. Next, we derived (see Supplementary Materials) the probability density function *P(f)* to observe an individual adhesion bond at a particular force (Fig. 3B), under the steady-state force loading and dissociation condition:

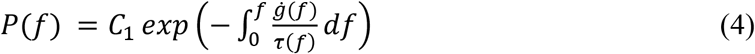

where *τ(f)* is the force-dependent dissociation of an adhesion complex, *g(f)* is the inverse function of *f(t), ġ(f)* is the derivative of *g(f)* with respect to force *f*, and *C*_*1*_ is the normalization constant. Given uniform surface densities of receptors on both the bead and substrate in our experiment settings, *P(f)* provides the generalized steady-state instantaneous force distribution. Given the force history of each adhesion bond, the probability density of the bond rupture force can be evaluated (Fig. 3D):

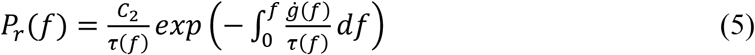

Where *C*_*2*_ is the normalization constant.

As the bead rolling velocity increases, the instantaneous force distribution quickly shifts up, capped by the unzipping of the reporting DNA at ∼15pN (Fig. 3B). The rupture force shows a bimodal distribution characteristic of catch-bond, with the distribution shifting towards the higher force mode as rolling velocity increases (Fig. 3D). The adhesion interaction between P-selectin and PSGL1 was described by a catch-bond, where the force-dependent bond life-time increases with force initially before it falls off again like a classic slip bond (*2, 8*). The DNA reporter in series with the P-selectin/PSGL1 bond modifies the overall adhesion characteristic, but only at forces above 13.6 pN, where the life-time of the DNA is lower than the P-selectin/PSGL1 interaction (Fig. 3D). At forces below 13.6 pN, the overall adhesion characteristic remains near identical to the P-selectin/PSGL1 interaction (Fig. 3D). Hence, in the adhesion footprint assay, the modified adhesion complex involving the DNA reporter remain a catch-bond up to 13.6 pN. Because of this catch-bond characteristic, both expected values of the instantaneous and rupture forces increase as the rolling velocity increases in a highly non-linear fashion (Fig. 3C). Below 13.6 pN, the life-time of P-selectin/PSGL1 is significantly shorter than the DNA reporter, making them statistically much more likely to rupture. Therefore, at low rolling velocity, the bond rupture event is predominantly P-selectin/PSGL1. Similarly, at high rolling velocity, the unzipping of DNA reporter dominates the overall bond rupture events (Fig. 3E). The greater fraction of DNA unzipping at high velocity correspond to the higher force peak ∼15 pN in the rupture force distribution (Fig. 3D), while the bond rupturing events happening < 5pN are due to P-selectin/PSGL1 dissociation.

Because the fluorescence intensity in the adhesion footprint assay is directly proportional to the amount of unzipped DNA reporters, it provides a direct link between experimental observables and model parameters. The tracks’ normalized fluorescence intensities as a function of their corresponding rolling velocity is in good agreement with our model (Fig. 3F). Therefore, through this model, a calibration curve can be established (Fig. 3G) to relate the fluorescence intensity along a track to the molecular forces the bead experiences at any given point (Fig. 3G).

### Quantitative mapping of molecular force along a single track

We isolated a single fluorescence track (Fig. 4B) to reconstruct the history of molecular adhesion force governing the rolling adhesion of a single bead. First, we straightened the track and removed the background fluorescence such that the fluorescence intensity is directly proportional to the amount of unzipped DNA reporters (Fig. 4C). Different segments along the track corresponding to different shear stress are clearly visible as bands in the straightened image (Fig. 4C). The transition points between two adjacent shear stress were determined by binary segmentation (*29*) of the mean intensity along the track (Fig. 4D). Determining the starting and end positions of each constant shear stress segment allows us to measure the mean fluorescence intensity and mean velocity of each segment (Fig. 4E), which can be fitted to our model with the diameter of the bead being the only fitting parameter. By doing so, the fluorescence-to-force calibration curve was established specifically for a single-bead track (Fig. 4F). Due to the size variation among the beads, it is necessary to determine this calibration curve for each single-bead track. The sizes of beads affect the reconstruction of molecular force because their different geometries lead to different force-loading profiles, even if they travel at the same linear velocity. Different force loading profile will subsequently lead to different instantaneous and rupture force distributions, and ultimately the molecular force reconstruction. From the model, the mean instantaneous force increases non-linearly with normalized fluorescence intensity (equivalent to the fraction of DNA rupture), while the mean rupture force scales nearly linearly (Fig. 4F). Hence, this force calibration curve allows us to reconstruct and map the mean and rupture force along the track (Fig. 4G, H).

**Fig. 4.**
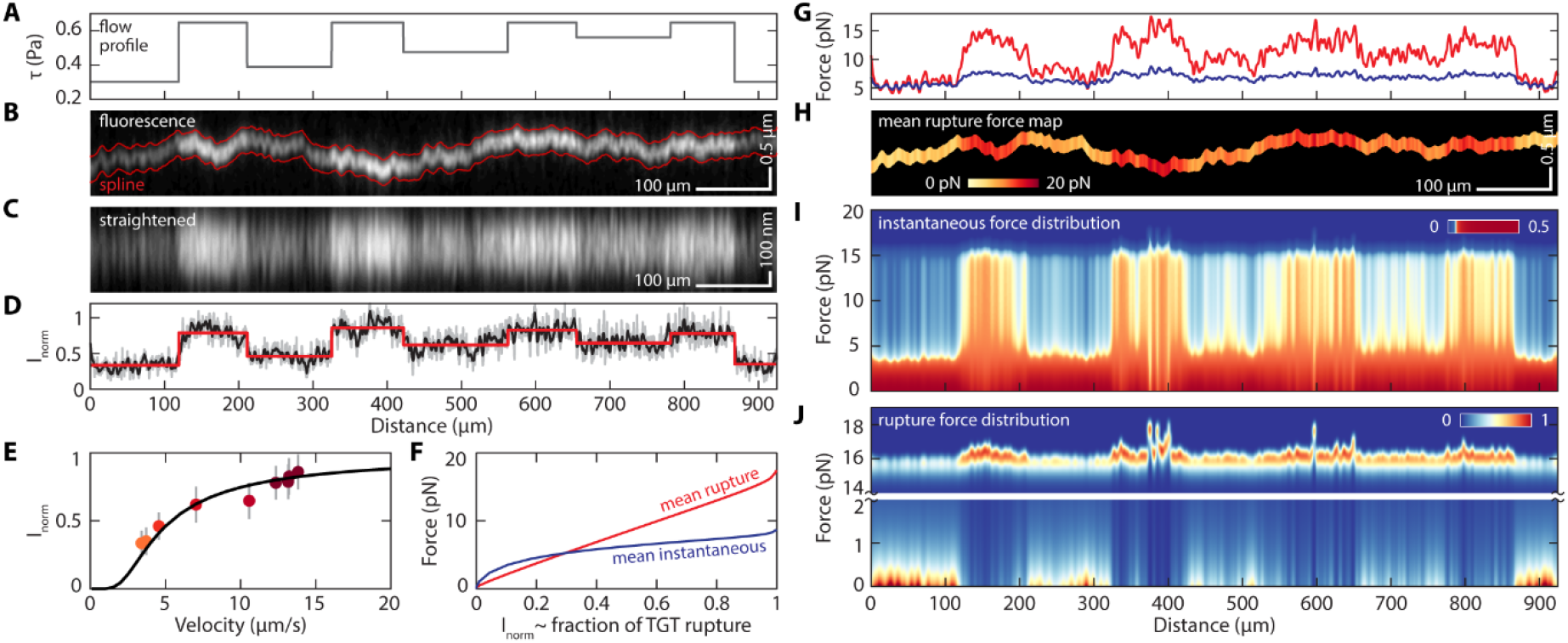
Mapping molecular force along a single rolling adhesion track. **(A)** Shear stress profile over time. **(B)** A single fluorescence track made by a bead rolling under this the flow profile shown in (A) where the spline is marked in red. **(C)** Straightened fluorescence track seen in (B) showing regions of high and low fluorescence intensity corresponding to the shear stress profile in (A). **(D)** The normalized total fluorescence intensity along the track (raw, gray; smoothed, black; fitted, red). **(E)** The normalized fluorescence intensity of each flow segment as a function of its mean bead rolling velocity. Data points are color-coded by shear stress, error bars indicate standard deviation of the fluorescence intensity in each segment; solid black line is the theoretical prediction of fluorescence intensity over bead velocity. **(F)** Calibration curve indicating mean instantaneous (blue) and rupture (red) force as a function of observed normalized fluorescence intensity (proportional to the fraction of ruptured TGT). **(G)** The mean instantaneous (blue) and rupture (red) force along the track. **(H)** Spatial mapping of the mean rupture force along the track. **(I)** The distribution of instantaneous force along the track. **(J)** The distribution of rupture force along the track. The image is split into a high force (14-19 pN) and a low force (0-2 pN) range.

While the mean instantaneous force is well defined, the mean rupture force should be interpreted more carefully, especially when a catch bond is involved. In contrast to a slip bond, the rupture force distribution is a bimodal distribution (Fig. 3D) instead of a unimodal distribution. Hence, changes of its expected value at different force loading rate comes mostly from the shift in the balance of its bimodal distribution. In contrast, slip bonds produces a unimodal rupture force distribution, and changes of expected value is more representative of a shift in its distribution. Because the instantaneous velocity of the bead correlates directly with the fluorescence intensity (Fig. 4E), and the molecular force distribution at any instantaneous velocity can be calculated from the model, we are able to construct the full molecular force distribution along the entire track (Fig. 4I). While forces below ∼4 pN is the dominant population throughout the entire track with various shear stress (0.3 -0.65 Pa), the force distribution in the 4-15 pN region is markedly more populated with increasing shear stress and rolling velocity (Fig. 4I). Similarly, under higher shear and velocity, the rupture force distribution moves from mainly <1 pN in the first, third, and last flow steps (Fig. 4J), to mostly between 15-18 pN, indicating a shift from mostly P-selectin/PSGL1 bond rupture (<1 pN) to mostly DNA unzipping (15-18 pN). This is precisely indicative of the overall catch-bond behaviour of the hybrid P-selectin/PSGL1 and DNA unzipping system.

## Discussion

Of the CAMs, there have been numerous studies and methods developed to characterize and monitor molecular forces on integrins (*15, 16, 30*–*32*), T-cell receptors (*17, 18, 33*), immunoglobulins (*34, 35*), cadherins (*19*–*21*), and others. However, the molecular forces of selectins in their physiological roles in rolling adhesion are the least understood. This is largely due to the unique challenges in observing selectin adhesion comparing to other families of CAMs. For example, α5β1 integrins are involved in initial attachment, firm adhesion, and motility of a cell on the extracellular matrix with high affinity, high receptor density, and relatively slow dynamics (*36, 37*). This is a system well-suited for reversible digital molecular force sensors (*16*). In the case of cadherins, the low receptor density allows individual receptor and force sensor to be imaged and is well suited for reversible analog force sensors (*21*). Unlike most CAMs, selectin adhesion during cell rolling is transient and highly dynamic as the cells travel at speed reaching many micrometers per second. In addition, each P-selectin/PSGL1 interaction only forms and ruptures once for durations on the order of millisecond. Therefore, it is not possible to obtain repeated measurements of the same interaction over time. This makes it infeasible to apply molecular force probes successfully used in other adhesion systems to the selectin adhesion during rolling adhesion. To support rolling adhesion, the surface receptor density needs to reach a minimum of ∼40 per µm^2^, much higher receptor density is required to support stable rolling. Hence this would require a high density of molecular force sensors on the surface, while at the same time, only ∼10-40 receptors per µm^2^ are engaged in the adhesion. Reversible sensors that require FRET or quenching to report force do not have sufficient signal-to-noise ratio to report at this level. In addition, it is practically challenging to catch a rolling bead in the field of view in real time, and the field of view of a 100x lens for TIRF imaging isn’t sufficiently large to allow us to follow a single bead over multiple shear stresses. Therefore, the only viable option to measure molecular forces in this system is using irreversible force sensors. It has high signal-to-noise ratio as the unruptured probes will not be fluorescently labelled, only probes subjected to force will rupture. Although not all molecular adhesion events produce a fluorescence signal, and this method will not measure the force history of a single receptor, the ensemble measurement allows us to extract precisely the distribution of molecular force. Our method allows for the extraction of quantitative force information from a record of force rupture events. The key to the success of this method is that we are measuring the fluorescence signal of a single bead across multiple shear stress. With the assistance of a model, this measurement allows us to make an individual calibration and subsequent quantification of molecular force. This is the first time that experimental force quantification was achieved using irreversible force sensors.

Contrary to previous observations where the cell rolling velocity decreases as the shear increases, which was attributed to a demonstration of the catch bond behaviour (*2*), our model and experiment shows that in a stable rolling model system, the rolling velocity is exponentially dependent on the shear stress. The mean and rupture molecular forces on individual adhesion tethers also increase monotonically to shear stress. Indeed, the previous observation could be observing a regime where the adhesion interaction is so sparse due to the low shear, that the apparent higher velocity of the cell is closely following the flow of the fluid. And in that system, the bead is not under stable rolling, but transient attachment. This apparent cell velocity is a hallmark of catch bond. In a system where rolling is stable due to a high density of surface adhesion interactions, even though the catch bond is working, it does not exhibit the type of behaviour reported previously.

In order to quantify tension using TGT-based force sensor, one must know the force-dependent lifetime of the receptor-ligand pair it is coupled to. The fluorescence signal reported by the TGT is determined by the ratio of rupture probability of the DNA against the ligand. For the particular TGT sequence in the unzipping geometry coupled to P-selectin/PSGL1, the crossover of dissociation lifetime occurs at ∼13.6 pN (as shown in Fig.3c). In other words, at forces below 13.6 pN the dissociation events are predominantly between P-selectin and PSGL1, while above 13.6 pN, the TGT rupture events dominate, generating fluorescent signal. Furthermore, around the crossover point, the changes of TGT lifetime as a function of force is significantly faster than that of the P-selectin/PSGL1interaction. This results in a sharp transition to primarily TGT rupturing and potentially limits the dynamic range of force sensing using TGT. Therefore, the choice of TGT sequence and rupture mode (i.e. unzipping or shearing) determines the molecular force range the assay is most sensitive to. To expand the dynamic range of TGT-based force sensors, a ratiometric approach using serially connected TGT can potentially be used (*38*).

In conclusion, we developed a new force quantification method specifically to address the challenges of studying molecular adhesion interactions that occur fast and only once. We combined the adhesion footprint assay and bead rolling as a model system to study interactions between P-selectin and PSGL1. The adhesion footprint allowed us to keep a fluorescent record of individual molecular adhesion events, where the fraction of ruptured TGTs is proportional to the observed fluorescence intensity. With the use of a model, we can use the intensity of the adhesion footprint to determine the molecular force distribution on the P-selectin/PSGL1 interactions during rolling. Our experimental results are in excellent agreement with our steady-state bead-rolling model. We found that with shear stresses increasing from 0.3 to 0.65 Pa, the fluorescence footprint intensity increased, corresponding to an increase of mean molecular rupture forces from ∼5 to 15 pN. This quantitative adhesion footprint assay addressed a fundamental challenge in quantifying brief molecular adhesion events and allowed us to study selectin-mediated interactions during the highly dynamic rolling adhesion. We believe that this assay can be generally applied to quantify the molecular forces involved in other fast molecular adhesion interactions.

## Materials and Methods

### Surface PEG passivation

To minimize non-specific adhesion of beads, the glass surfaces of the coverslip and top slide were passivated with polyethylene glycol (PEG) following previously established protocol (*27*). Briefly, this was accomplished by first submerging the coverslips and slides in a piranha solution (3:1 sulfuric acid to hydrogen peroxide) for 30 minutes and then copiously rinsing first with milli-Q water and then with methanol. Following the methanol rinse, the coverslips/slides were placed in a 1% aminosilane solution (94 mL methanol, 1 mL 3-(2-aminoethylamino)propyltrimethoxysilane (VWR, CAS# 1760-24-3), 5 mL glacial acetic acid) for 1 hour at 70°C. Once the silanization was complete, a methanol rinse and then a water rinse was done before putting the coverslips/slides in the oven. The coverslips/slides were placed in an oven at 110°C for 20 minutes and then left to cool to room temperature. Once at room temperature the coverslips/slides were passivated with PEG (Laysan Bio, mPEG-SVA, 5k) and/or biotinylated PEG (Laysan Bio, biotin-PEG-SVA, 5k) by placing 80 µL of 250 mg/mL PEG solution on one surface of the coverslips/slides and then placing a second coverslip/slide on-top of the PEG. The PEG passivation was left to react overnight at room temperature, and then the passivated coverslips/slides were washed with milli-Q water, dried with nitrogen and stored at -20°C under nitrogen until they were needed for an experiment. Each time coverslips/slides were PEGylated, 8 coverslips/slides were passivated and stored for future experiments.

### Parallel plate flow chamber

For each bead rolling experiment, a parallel plate flow chamber was constructed with a coverslip (Fisherbrand Premium, 12-548-5P), permanent double-sided tape (Scotch 3M237), and a top microscope slide (VWR, 48300-026). Prior to construction of the flow chamber, the coverslip was passivated with 20:1 PEG to PEG-biotin and the top slide was passivated with PEG. To create inlet and outlets for the flow chamber, a Dremel was used to drill holes on either end of the top slide. Channels were cut into the double-sided tape with a laser engraver (BOSS LASER) and then the tape was sandwiched between the coverslip and the top slide to create channels with dimensions 0.093 x 53.5 x 2.2mm. The flow chamber was then mounted on a custom-made bracket with inlet and outlet adapters allowing for a syringe pump to be used with the chamber. The syringe pump was used to apply laminar flow through the chamber at variable shear stresses between 0.30 and 0.65 Pa with a specific, user determined, flow series.

### PSGL1 bead preparation

Protein G-coated polystyrene beads (Spherotech, PGP-60-5) were functionalized with PSGL1-Fc Chimera (R&D systems, 3345-PS-050) as described previously (*19*). 10 µL of stock beads (0.5% w/v) were washed twice with PBS (pH 7.4). To wash the beads, they were centrifuged at 3000 g for 15 minutes, the supernatant was discarded, and the beads were resuspended in PBS. Following the second resuspension in PBS, the beads were spun down a third time and resuspended in 50 µL of 100 µg/mL PSGL1-Fc Chimera and left to incubate in the PSGL1 solution for >2 hours on a rotator at room temperature. Prior to each experiment, the beads were washed with T50M5C2 (10 mM Tris, 50 mM NaCl, 5 mM MgCl_2_ and 2 mM CaCl_2_) to remove excess PSGL1 and prepare for rolling.

### Preparation of TGT for adhesion footprint assay

The TGT DNA construct used for the bead rolling adhesion footprint assay was based on previously described protocols (*27*). The assay involved a biotinylated bottom strand, a protein G functionalized top strand, a fluorescently labeled probe strand, and a blocker top strand. The following oligonucleotides and modifications were ordered from Integrated DNA Technologies (IDT):

Biotin bottom stand: 5’-/5BiotinTEG/ TTTTT CCCTCCTGCGTCGCCCGG-3’

Thiol top strand: 5’-CCGGGCGACGCAGGAGGG TTTTT /3ThioMC3-D/-3’

Blocker top strand: 5’-CCGGGCGACGCAGGAGGG-3’

Probe strand: 5’-CCGGGCGACGCAGG /3Cy3Sp/-3’

The top strand was conjugated to His-tagged Protein G (Abcam, ab49807) through a sulfo-SMCC (Thermofisher, 22322) hetero-bifunctional crosslinker that linked the 3’ thiol on the top strand to an amine moiety of the Protein G. The conjugation was performed following the protocols described by the sulfo-SMCC manufacturer. Following conjugation, the product was purified using His-Tag Isolation and Pulldown Dynabeads. The conjugated product (Top-PG) was hybridized to the bottom strand by mixing the oligos at a molar ratio of 2:1 (400:200 nM) in T50M5C2 buffer ensuring an excess of Top-PG. The hybridization was performed at room temperature for 2 hours prior to experiments resulting in the full biotin and protein G functionalized TGT (TGT-biot-PG).

### Adhesion footprint assay

The parallel plate flow chamber was functionalized by incubating BSA (TOCRIS Bioscience, 9048-46-8), streptavidin (Cedarlane, CL1005-01-5MG), TGT-biot-PG and P-selectin Fc chimera (R&D Systems, 137-PS-050). BSA (10 mg/mL) was flowed into the chamber and incubated for 15 minutes to further passivate the PEGylated surface ensuring minimal non-specific interaction with the glass substrate. The excess BSA was washed out of the chamber with 200 µL T50M5C2 buffer at a flow rate of 2 mL/hr controlled by a syringe pump (Harvard apparatus). All wash steps were performed in this manner. Streptavidin (100 µg/mL) was then flowed into the chamber and incubated for 15 minutes to bind to the biotin on the PEG/PEG-biotin surface. The streptavidin was washed out with T50M5C2 and then TGT-biot-PG (200 nM) was flowed into the chamber and incubated for 15 minutes allowing the streptavidin on the surface to bind to the biotin on the TGT-biot-PG. Excess TGT-biot-PG was washed out with T50M5C2 and then P-selectin Fc chimera (10 µg/mL) was incubated for 15 minutes allowing the Protein G to bind to the immunoglobulin G (IgG) Fc domain of the P-selectin Fc chimera. Excess P-selectin was then washed out with T50M5C2. Finally, blocker top strand DNA (200 nM) was incubated in the chamber for 15 minutes to hybridize to any remaining “empty” bottom strand DNA on the surface that failed to hybridize to the Top-PG. This step ensured that there was minimal ssDNA on the surface of the chamber.

Once the chamber was functionalized with P-selectin and TGT, the PSGL1 beads were flowed into the chamber and left to settle for 5 minutes. A syringe pump was then used to apply a specific flow series of shear stresses in the range of 0.30 and 0.65 Pa causing the PSGL1 beads to roll on the P-selectin/TGT surface. The rolling beads were observed on a home-made darkfield microscope with a 10X objective and the images were recorded at 30 fps. Following bead rolling, the probe strand (100 nM) was flowed into the chamber in T50M5C2 buffer and incubated for 5 minutes to label the ruptured TGTs allowing for observation of the fluorescence tracks left by the rolling beads. The excess probe strand was washed out of the chamber with imaging buffer (40 mM NaCl, 160 mM Tris, 10% (m/v) Glucose, 1.12 mg/mL Glucose Oxidase (Sigma, G7141-50KU), 0.08 mg/mL catalase (Sigma, C9322), pH 8.0) prior to fluorescence imaging.

### TIRF imaging and fluorescence image processing

The sample was excited with a 532nm laser (Spectra-Physics, Excelsior 532) and observed through TIRF microscopy on an Olympus IX83 inverted microscope. The fluorescence images were acquired with an Andor iXon Ultra 897 EMCCD camera. Because the length of fluorescence track from each bead is on the order of mm, while the TIRF field-of-view is only ∼80×80 µm^2^, ∼2000-3000 individual images were acquired with the motorized XY stage to produce a large stitched image. The microscope was programmed to move so that there was a 10% overlap between adjacent images for image registration during tiling. Following image acquisition, additional processing was done to correct for uneven illumination in the original images (*I*_*0*_) following the protocol described previously (*27*). During each experiment, a background image (*I*_*bg*_) was acquired by taking an image with the illumination turned off. This was used to do a background subtraction (*I*_*0*_ *– I*_*bg*_) to obtain the true fluorescence signal of each image. To determine the underlying illumination profile (*I*_*illumination*_), the average of all the background subtracted images *<I*_*0*_ *– I*_*bg*_*>*, was calculated and normalized its maximum to 1. The flattened illumination profile (*I*_*flat*_) was calculated using the formula: *I*_*flat*_ *= (I*_*0*_ *– I*_*bg*_*)/I*_*illumination*_. The flattened images were then used to construct the final stitched image using the ImageJ Grid/Collection stitching plugin with linear blending selected as the chosen fusion method.

### Fluorescence track processing and analysis

Fluorescence track processing and analysis was performed on the stitched image through the following steps: track detection, tracking, isolation, and analysis. Track detection was done to locate the pixels corresponding to fluorescence tracks. This was accomplished by finding the peak location of each column in the image. The detected peaks were then representative of track cross-sections and the position of the peak maxima was recorded as a point belonging to a fluorescence track. During detection, thresholds were applied to only detected peaks that were ∼1.5 track widths away from each other. Track width was manually determined by counting the number of pixels spanning the cross section of a track. This peak proximity threshold ensured that the fluorescence intensities were not influence by neighbouring tracks. Upon acquiring the pixel coordinates belonging to the fluorescence tracks, custom-written MATLAB code was used to assign coordinates to tracks and sequentially append each coordinate to its corresponding track. Track assignment was determined based on the proximity of a coordinate to the end position of the neighboring tracks. Each track was then cropped out of the stitched image by extracting the track coordinates ± half the track width for the full length of the track.

Fluorescence intensity traces of each track was then done by calculating the mean intensity of each column along the length of the track. Once the intensity traces were obtained for each track, they were subdivided into segments corresponding to the flow rates applied during the rolling assay. This segmentation was done by determining abrupt changes in the mean of a dataset. This function was sensitive enough to detect abrupt changes in the fluorescence intensity along the length of the tracks corresponding to changes in flow rate. The accuracy of the segmentation was verified by a comparison with the segments in the applied flow series. The detected segments were then used to determine the average intensity and the bead velocity corresponding to the applied flow rates.

### Bead detection and tracking

Bead rolling velocity was extracted from the 10x darkfield images with custom-written MATLAB code that detects and tracks the position of individual beads over time. Bead detection was achieved using the built-in MATLAB function, *imfindcircles*, that uses a circular Hough transform to find circles in an image. This function works exceptionally well with beads as they appear to be perfect circles under the microscope. Once the beads were detected, the centroid position of each bead was recorded on a frame-by-frame basis. In a similar fashion to how the fluorescence tracks were tracked, for every frame, each bead was assigned to a track. This allowed for the analysis of each bead’s displacement on a frame-by-frame basis, and hence, the calculation of the bead instantaneous velocity. With this approach, we were able to calculate the observed bead rolling velocity and use it as a standard to compare the fluorescence track velocities with.

### Super-resolution Adhesion Footprint

To acquire a super-resolution image of the bead tracks, the probe strand used for diffraction limited images was replaced with the following sequence purchased from IDT:

DNA PAINT imager strand: 5’-GAGGGAAATT/3Cy3Sp/-3’

The DNA PAINT imager strand was designed following literature recommendations (*39*). Immediately following bead rolling, 500pM of DNA PAINT imager strand in DNA PAINT buffer (0.05% Tween-20, 75 mM MgCl_2_, 5 mM Tris, and 1 mM EDTA) was added to the chamber. Upon adding the imager strand, the sample was observed through TIRF with the same microscope, camera and laser used to image the diffraction-limited fluorescence tracks. The DNA PAINT “blinking” was imaged for 50,000 frames at an exposure time of 25ms. Upon acquiring the fluorescence images, the ‘Picasso’ software package (*40*) was used to identify the position of all the fluorophores in every frame and then construct the super-resolution image. The final super-resolution image was then drift corrected with ‘Picasso’ through redundant cross-correlation.

### Statistical analysis

MATLAB was used to calculate the Pearson correlation coefficient.

## H2: Supplementary Materials

Note S1. Derivation of the steady state model.

Fig. S1. Bead rolling velocity scales exponentially to shear stress.

## General

we would like to thank Yousif Murad for discussions.

## Funding

we thank the Natural Sciences and Engineering Research Council of Canada (NSERC RGPIN-2017-04407), Canada Foundation for Innovation (CFI 35492), and the Michael Smith Foundation for Health Research (Scholar Award) for support.

## Author contributions

AY and IL designed the experiment, AY performed the experiment and analyzed data, IL developed the model, AY and IL drafted the manuscript.

## Competing interests

none.

## Data and materials availability

all data available upon request.

## Notes

### Competing Interest Statement

The authors have declared no competing interest.

## References

1. S. Nourshargh, R. Alon, Leukocyte Migration into Inflamed Tissues. Immunity. 41 (2014), pp. 694–707.

2. R. P. McEver, C. Zhu, Rolling cell adhesion. Annu. Rev. Cell Dev. Biol. 26 (2010), pp. 363–396.

3. A. Zarbock, K. Ley, Neutrophil Adhesion and Activation under Flow. Microcirculation. 16, 31–42 (2009).

4. J. Fritz, A. G. Katopodis, F. Kolbinger, D. Anselmetti, Force-mediated kinetics of single P-selectin/ligand complexes observed by atomic force microscopy. Proc. Natl. Acad. Sci. U. S. A. 95, 12283–12288 (1998).

5. J. A. Simmons, W. J. Bailey, M. D. Greenfield, T. E. Shelly, E. Coscia, P. D. Phillips, J. C. Fentress, Lifetime of the p-selectin-carbohydrate bond and its response to tensile force in hydrodynamic flow. Nature. 374 (1995), pp. 539–542.

6. P. Mehta, R. D. Cummings, R. P. McEver, Affinity and kinetic analysis of P-selectin binding to P-selectin glycoprotein ligand-1. J. Biol. Chem. 273, 32506–32513 (1998).

7. B. T. Marshall, M. Long, J. W. Piper, T. Yago, R. P. McEver, C. Zhu, Direct observation of catch bonds involving cell-adhesion molecules. Nature. 423, 190–193 (2003).

8. M. T. Beste, D. A. Hammer, Selectin catch-slip kinetics encode shear threshold adhesive behavior of rolling leukocytes. Proc. Natl. Acad. Sci. U. S. A. 105, 20716–20721 (2008).

9. H. B. Wang, J. T. Wang, L. Zhang, Z. H. Geng, W. L. Xu, T. Xu, Y. Huo, X. Zhu, E. F. Plow, M. Chen, J. G. Geng, P-selectin primes leukocyte integrin activation during inflammation. Nat. Immunol. 8, 882–892 (2007).

10. Y. Zhang, G. Sun, S. Lü, N. Li, M. Long, Low spring constant regulates P-selectin-PSGL-1 bond rupture. Biophys. J. 95, 5439–5448 (2008).

11. W. Hanley, O. McCarty, S. Jadhav, Y. Tseng, D. Wirtz, K. Konstantopoulos, Single molecule characterization of P-selectin/ligand binding. J. Biol. Chem. 278, 10556–10561 (2003).

12. S. Lü, Z. Ye, C. Zhu, M. Long, Quantifying the effects of contact duration, loading rate, and approach velocity on P-selectin-PSGL-1 interactions using AFM. Polymer (Guildf). 47, 2539–2547 (2006).

13. E. Evans, A. Leung, V. Heinrich, C. Zhu, Mechanical switching and coupling between two dissociation pathways in a P-selectin adhesion bond. Proc. Natl. Acad. Sci. U. S. A. 101, 11281–11286 (2004).

14. L. J. Rinko, M. B. Lawrence, W. H. Guilford, The Molecular Mechanics of P-and L-Selectin Lectin Domains Binding to PSGL-1. Biophys. J. 86, 544–554 (2004).

15. M. Morimatsu, A. H. Mekhdjian, A. S. Adhikari, A. R. Dunn, Molecular tension sensors report forces generated by single integrin molecules in living cells. Nano Lett. 13, 3985–3989 (2013).

16. Y. Zhang, C. Ge, C. Zhu, K. Salaita, DNA-based digital tension probes reveal integrin forces during early cell adhesion. Nat. Commun. 5, 1–10 (2014).

17. Y. Liu, L. Blanchfield, V. Pui-Yan Ma, R. Andargachew, K. Galior, Z. Liu, B. Evavold, K. Salaita, DNA-based nanoparticle tension sensors reveal that T-cell receptors transmit defined pN forces to their antigens for enhanced fidelity. Proc. Natl. Acad. Sci. U. S. A. 113, 5610–5615 (2016).

18. V. P. Y. Ma, Y. Liu, L. Blanchfield, H. Su, B. D. Evavold, K. Salaita, Ratiometric tension probes for mapping receptor forces and clustering at intermembrane junctions. Nano Lett. 16, 4552–4559 (2016).

19. X. Wang, Z. Rahil, I. T. S. Li, F. Chowdhury, D. E. Leckband, Y. R. Chemla, T. Ha, Constructing modular and universal single molecule tension sensor using protein G to study mechano-sensitive receptors. Sci. Rep. 6, 1–10 (2016).

20. D. Eder, K. Basler, C. M. Aegerter, Challenging FRET-based E-Cadherin force measurements in Drosophila. Sci. Rep. 7, 1–12 (2017).

21. D. E. Conway, M. T. Breckenridge, E. Hinde, E. Gratton, C. S. Chen, M. A. Schwartz, Fluid shear stress on endothelial cells modulates mechanical tension across VE-cadherin and PECAM-1. Curr. Biol. 23, 1024–1030 (2013).

22. F. Meng, T. M. Suchyna, E. Lazakovitch, R. M. Gronostajski, F. Sachs, Real time FRET based detection of mechanical stress in cytoskeletal and extracellular matrix proteins. Cell. Mol. Bioeng. 4, 148–159 (2011).

23. S. K. Bhatia, M. R. King, D. A. Hammer, The state diagram for cell adhesion mediated by two receptors. Biophys. J. 84, 2671–2690 (2003).

24. E. Evans, A. Leung, D. Hammer, S. Simon, Chemically distinct transition states govern rapid dissociation of single L-selectin bonds under force. Proc. Natl. Acad. Sci. U. S. A. 98, 3784–3789 (2001).

25. H. Läubli, L. Borsig, Altered Cell Adhesion and Glycosylation Promote Cancer Immune Suppression and Metastasis. Front. Immunol. 10 (2019), p. 2120.

26. A. Yasunaga, Y. Murad, I. T. S. Li, Quantifying molecular tension-classifications, interpretations and limitations of force sensors. Phys. Biol. 17 (2020), p. 011001.

27. I. T. S. Li, T. Ha, Y. R. Chemla, Mapping cell surface adhesion by rotation tracking and adhesion footprinting. Sci. Rep. 7, 1–11 (2017).

28. X. Wang, T. Ha, Defining single molecular forces required to activate integrin and Notch signaling. Science (80-.). 340, 991–994 (2013).

29. R. Killick, P. Fearnhead, I. A. Eckley, Optimal detection of changepoints with a linear computational cost. J. Am. Stat. Assoc. 107, 1590–1598 (2012).

30. D. R. Stabley, C. Jurchenko, S. S. Marshall, K. S. Salaita, Visualizing mechanical tension across membrane receptors with a fluorescent sensor. Nat. Methods. 9, 64–67 (2012).

31. Y. Liu, R. Medda, Z. Liu, K. Galior, K. Yehl, J. P. Spatz, E. A. Cavalcanti-Adam, K. Salaita, Nanoparticle tension probes patterned at the nanoscale: Impact of integrin clustering on force transmission. Nano Lett. 14, 5539–5546 (2014).

32. F. Li, S. D. Redick, H. P. Erickson, V. T. Moy, Force measurements of the α5β1 integrin-fibronectin interaction. Biophys. J. 84, 1252–1262 (2003).

33. K. H. Hu, M. J. Butte, T cell activation requires force generation. J. Cell Biol. 213, 535–542 (2016).

34. Z. Wan, X. Chen, H. Chen, Q. Ji, Y. Chen, J. Wang, Y. Cao, F. Wang, J. Lou, Z. Tang, W. Liu, The activation of IgM-or isotype-switched IgG-and IgE-BCR exhibits distinct mechanical force sensitivity and threshold. Elife. 4 (2015), doi:10.7554/eLife.06925.

35. N. Yeow, R. F. Tabor, G. Garnier, Mapping the distribution of specific antibody interaction forces on individual red blood cells. Sci. Rep. 7, 1–7 (2017).

36. R. Pankov, E. Cukierman, B. Z. Katz, K. Matsumoto, D. C. Lin, S. Lin, C. Hahn, K. M. Yamada, Integrin dynamics and matrix assembly: Tensin-dependent translocation of α5β1 integrins promotes early fibronectin fibrillogenesis. J. Cell Biol. 148, 1075–1090 (2000).

37. J. Takagi, K. Strokovich, T. A. Springer, T. Walz, Structure of integrin α5β1 in complex with fibronectin. EMBO J. 22, 4607–4615 (2003).

38. Y. Murad, I. T. S. Li, Quantifying Molecular Forces with Serially Connected Force Sensors. Biophys. J. 116, 1282–1291 (2019).

39. M. Dai, in Methods in Molecular Biology (Humana Press Inc., 2017; https://pubmed.ncbi.nlm.nih.gov/27813009/), vol. 1500, pp. 185–202.

40. J. Schnitzbauer, M. T. Strauss, T. Schlichthaerle, F. Schueder, R. Jungmann, Super-resolution microscopy with DNA-PAINT. Nat. Protoc. 12, 1198–1228 (2017).

